# (Epi)mutation rates and the evolution of composite trait architectures

**DOI:** 10.1101/2022.12.23.521798

**Authors:** Bastien Polizzi, Vincent Calvez, Sylvain Charlat, Etienne Rajon

**Affiliations:** Laboratoire de Mathématiques de Besançon, Université de Bourgogne Franche-Comté, Centre National de la Recherche Scientifique, UMR 6623, Besançon, France; Institut Camille Jordan, Université de Lyon, Université Claude Bernard Lyon 1, Centre National de la Recherche Scientifique, UMR 5208, Villeurbanne, France; Equipe-Projet Inria Dracula, Centre Inria de Lyon, France; Laboratoire de Biométrie et Biologie Evolutive, Université de Lyon, Université Claude Bernard Lyon 1, Centre National de la Recherche Scientifique, UMR 5558, Villeurbanne, France

## Abstract

Mutation rates vary widely along genomes and across inheritance systems. This suggests that complex traits – resulting from the contributions of multiple determinants – might be composite in terms of the underlying mutation rates. Here we investigate through mathematical modeling whether such an heterogeneity may drive changes in a trait’s architecture, especially in fluctuating environments where phenotypic instability can be beneficial. We first identify a convexity principle, related to the shape of the trait’s fitness function, setting conditions under which composite architectures should be adaptive or, conversely and more commonly, should be selected against. Simulations reveal, however, that applying this principle to realistic evolving populations requires taking into account pervasive epistatic interactions that take place in the system. Indeed, the fate of a mutation affecting the architecture depends on the (epi)genetic background, itself depending upon the current architecture in the population. We tackle this problem by borrowing the adaptive dynamics framework from evolutionary ecology – where it is routinely used to deal with such resident/mutant dependencies – and find that the principle excluding composite architectures generally prevails. Yet, the predicted evolutionary trajectories will typically depend on the initial architecture, possibly resulting in historical contingencies. Finally, by relaxing the large population size assumption, we unexpectedly find that not only the strength of selection on a trait’s architecture, but also its direction, depend on population size, revealing a new occurrence of the recently coined phenomenon of ‘sign inversion’.

## 1 Introduction

Complex traits, those resulting from the combined contributions of multiple determinants, may in principle be subject to composite mutations rates. For example, genes contributing to the same trait may mutate at rates spanning orders of magnitude because their genomic environments differ, or simply their lengths (Rando and Verstrepen 2007; Hodgkinson and Eyre-Walker 2011; Oman, Alam, and Ness 2022). The possibility of heritable yet non-genetic contributions introduces even more pronounced heterogeneity in mutation rates *sensu lato*, that is, in the rates of random change, whatever the inheritance mechanism (Rando and Verstrepen 2007; Danchin 2013). Indeed, non-genetic inheritance systems, as multiple as they may be, share the property of being less stable than DNA sequences (Johannes et al. 2009; Rechavi, Minevich, and Hobert 2011; Denkena, Johannes, and Colomé-Tatché 2021; Dodson and Rine 2015; Graaf et al. 2015). Here we investigate whether such an heterogeneity in mutation rates may drive the evolution of a trait’s architecture: just as mutation rates can be adaptively tuned by the degree of environmental stability (Ishii et al. 1989; Johnson 1999; Andre and Godelle 2006), can trait architectures evolve toward optimally mixed contributions of determinants differing in their rates of mutations? Or, on the contrary, are non-composite mutation rates generally selected for, which should lead to simplified traits architectures, homogeneous in terms of the underlying mutation rates?

We address these questions through mathematical modeling, assuming that selection acts on a single trait in an environment oscillating between two states. We further assume that the relative contributions of two (or more) determinants to the trait, differing only in their respective mutation rates, are controlled by one evolving parameter, and we study its evolution. In our analysis, the mutation rates differ by orders of magnitude, such that they can be referred to as ‘fast’ and ‘slow’ in the case of two determinants. We start with the analytical treatment of an idealized situation where only the absolute growth rate of a mutant determines its fate, and find the architectures that maximize this fitness proxy. We thereby identify a general mathematical principle setting boundaries to the conditions where complex architectures, and thus composite mutation rates, may be optimal. According to this principle, whatever the degree of environmental instability, non-composite mutation rates are favored as long as individuals of intermediate phenotypic value do not perform better across different environments than the average of the extreme phenotypes. In other words, if the fitness function of the trait is convex, favoring specialists over generalists, the fitness function of the trait architecture is also convex such that non-mixed contributions are selectively favored. However, we also show that, as long as the degree of environmental instability is not too high, non-composite mutation rates remain optimal even if the fitness function of the trait is concave.

Absolute growth rates, however, ignore a potentially important aspect of the evolutionary dynamics at play in this system: a mutation necessarily occurs in an (epi)genetic background whose state (i) may depend on the strategy in place, and (ii) may critically affect the mutant’s fate. We address this issue by using the adaptive dynamics framework, initially built to deal with ecological interactions by accounting explicitly for context-dependencies in a mutant’s invasion success (Metz, Nisbet, and Geritz 1992; Geritz et al. 1998). This approach generally confirms the above outlined convexity argument, but also reveals that pervasive epistatic interactions render the system reluctant to change, because determinants contributing little to the trait tend to accumulate cryptic variation that becomes deleterious if the trait architecture is changed. This reveals that historical contingencies may prove important in the evolution of genetic architectures.

Finally, we use simulations to relax the assumption of a large population size and thus investigate the consequences of genetic drift on the evolution of a trait’s architecture. In doing so, we reveal a counterintuitive pattern whereby reducing population size below a threshold changes not only the efficiency but also the direction of selection. We relate this finding to a recently described phenomenon, coined ‘sign-inversion’, that generally takes place whenever some source of variability produces selective pressures differing in their direction, average duration, and strength (Raynes, Wylie, et al. 2018; Raynes, Burch, and Weinreich 2021). In such situations, weak but stable selection in one direction is dominant in large populations, while strong but occasional selection in the other direction becomes the main player in small populations. In our case, sudden environmental changes strongly select for a large contribution of the fastest determinant whereas environment stasis selects for a reduced input of mutations and therefore a high contribution of the slowest determinant. Yet, the latter selection pressure is typically weak and long-lasting, thereby acting efficiently in large populations only, resulting in a trait’s architecture that is contingent on population size.

## 2 Models and tools

### 2.1 Qualitative summary of the model

#### Trait architecture

The model considers *I* ≥ 2 determinants with distinct mutation rates that contribute to a phenotypic trait, varying continuously from 0 to 1. Each determinant is characterized by (i) its (epi)genotype, taking a binary value (*X*_*i*_ = 0 or *X*_*i*_ = 1), (ii) its mutation rate (*μ*_*i*_ *>* 0) at which the (epi)genotype switches randomly from *X*_*i*_ to 1 − *X*_*i*_, and (iii) its relative contribution to the trait (*α*_*i*_ ∈ [0, 1]). The contributions sum to one, that is ∑_*i*_ *α*_*i*_ = 1. The resulting trait for a given individual is the weighted sum of its determinant values: Φ_*X*_(*α*) = ∑_*i*_ *α*_*i*_*X*_*i*_ (simply denoted as Φ if unambiguous).

In the following, the vector of contributions ***α*** = (*α*_*i*_)_1≤*i*≤*I*_ of a single individual is referred to as its *trait architecture*. In the simulations, we assume that the values of ***α*** are subject to mutational changes, at the slowest rate min *μ*_*i*_. Mutated ***α*** values are drawn in a Gaussian law centered on the parental value with variance 0.1, and constrained to satisfy ∑_*i*_ *α*_*i*_ = 1.

#### Notations in the case of two determinants (fast versus slow)

The main part of our study is devoted to the case of two determinants, below referred to as ‘fast’ and ‘slow’, in reference to their contrasted mutation rates *μ*_*F*_ and *μ*_*S*_ (*μ*_*F*_ ≫ *μ*_*S*_). In this particular case, we can denote the respective contributions of the fast and slow inheritance systems as *α*_*F*_ and *α*_*S*_ = 1 − *α*_*F*_. The (epi)genotype values are then denoted *X*_*F*_ and *X*_*S*_, both in {0, 1}.

#### Evolutionary dynamics

We use a birth and death model, continuous in time, so that generations are overlapping. The fecundity rate is assumed to be homogeneous. Up to changing units of time, it is set to 1. Selection acts on the rate of mortality, namely *s*|Φ − Φ^*^(*t*)|^*γ*^, where the strength of selection is controlled by parameter *s*, Φ^*^ is the environment-dependent optimal phenotype, and the exponent *γ* ≥ 0 is the shape parameter of the fitness function (see Figure 1). We distinguish three cases on the basis of *γ* values: *γ* = 1 is our reference case, and corresponds to a linear relationship between trait and fitness; *γ <* 1 corresponds to a convex relationship resulting in a higher average fitness of *specialists* (Φ close to extremal values 0 or 1); in contrast *γ >* 1 results in a concave relationship favoring *generalists* (Φ close to 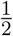).

**Figure 1:**
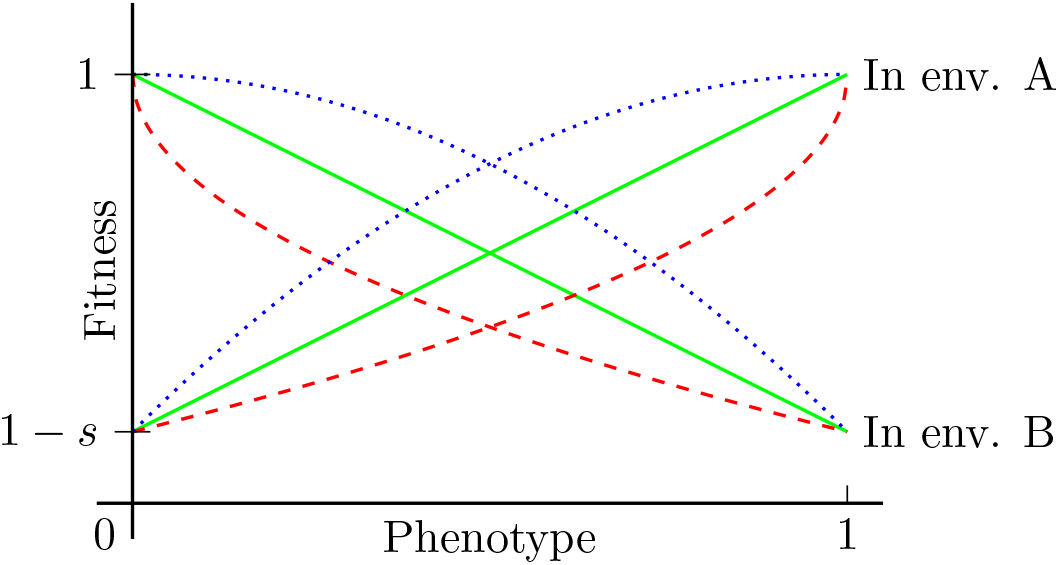
Shape of the fitness function. 1 − *s*|Φ − Φ^*^|^*γ*^, for *γ* = 1 (plain green, linear case), *γ* = 1*/*2 (dashed plain red, strictly convex case) and *γ* = 2 (dotted blue, strictly concave case).

We assume that population size is regulated by an additional density-dependent contribution to the mortality rate 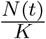, where *N* (*t*) is the instantaneous population size, and *K* denotes the carrying capacity.

#### Environmental variation

We assume that the environment fluctuates between two possible states 𝒜/ℬ associated with different optimal phenotypes 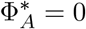 and 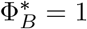. We consider periodic changes of period *T*, with a symmetrical alternance between 𝒜 and ℬ. It is relevant to distinguish three regimes of environmental variation in relation with mutation rates. First, the environment may be changing very rarely, that is, even more slowly than the slowest determinant (*T* (min *μ*_*i*_) ≫ 1). Second, the environment may be changing very often, that is much faster than the least stable determinant (*T* (max *μ*_*i*_) ≪ 1). A third, intermediate regime, is one where the rate of environmental change is comparable to the range of mutation rates.

#### Previous occurrence of the model

The model is inspired by the study in (Rajon and Charlat 2019) who restricted their model to the case of two determinants (*I* = 2), with respective contributions *α*_1_ = 1 − *α* and *α*_2_ = *α*, and to *γ* = 1. In their interpretation, the number *α* ∈ [0, 1] measures the balance between genetic and epigenetic contributions to the trait value. The only difference lies in the mutation rate, being significantly larger for the epigenetic contribution. While not limiting our interest to epigenetic inheritance, we retain from this initial work the focus on the variability of the mutation rates *μ*_*i*_’s and its consequences on the evolution of *α*. Separation of mutation timescales remains indeed the main component of our mathematical analysis.

### 2.2 A Lyapunov exponent approach in infinite populations

We begin our analysis by considering a monomorphic population for the trait architecture ***α*** = (*α*_*i*_)_1≤*i*≤*I*_, taking its growth rate as a proxy for its fitness. To this end, we introduce the Lyapunov exponent *λ*(***α***), which is the overall growth rate of the population over a complete cycle of environmental fluctuations. The rate *λ*(***α***) is measured in the long run, that is, once the respective frequencies of the (epi)genotypes (*X*_*i*_)_1≤*i*≤*I*_ have reached their periodic equilibrium distribution *f*_*g*_(*t*), where *g* denotes the (epi)genotype, taking value in 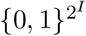. Technically, this equilibrium distribution is characterized by a periodic eigenvalue problem (Floquet spectral theory, see Appendix A), with the eigenvalue *λ*(***α***) defined by the following formula:

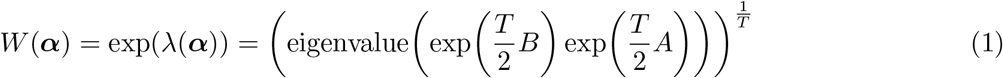

where *A* (resp. *B*) is the matrix describing the population dynamics in environment 𝒜 (resp. ℬ), which is defined by the various rates of mutation and mortality in the population. For instance, when *I* = 2, there are 2^2^ = 4 possible states in the population, namely (0, 0), (1, 0), (0, 1), (1, 1). In particular, the variation of the state (0, 1) results from: (i) mutations from the states (0, 0) and (1, 1) at respective rates *μ*_2_ and *μ*_1_, (ii) mutations to the states (0, 0) and (1, 1) at the same rates, (iii) birth-death dynamics specific to the state (0, 1) at rate 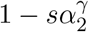.

The birth-death-mutation dynamics of the four states are then summarized in the following matrix

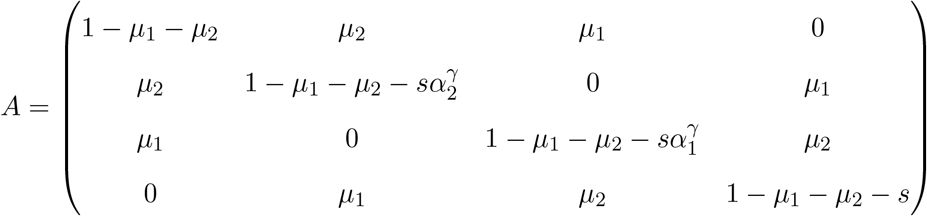

Note that the matrix *B* is the same, up to a symmetrical change in the diagonal:

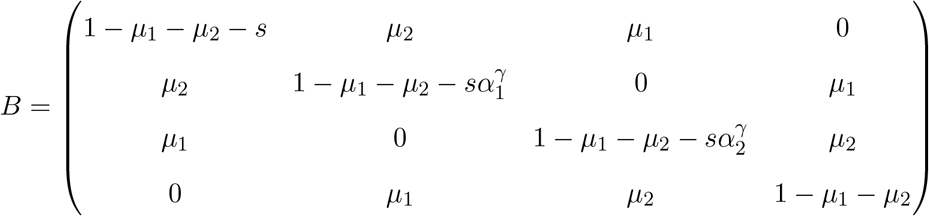

### 2.3 Adaptive dynamics

The Lyapunov exponent approach described above ignores the influence that the state of the population in place (denoted the ‘resident’, with trait architecture ***α***_*r*_), may have on the population dynamics of a ‘mutant’ with a different architecture ***α***_*m*_. The adaptive dynamics framework allows to take these interactions into account by calculating the mutant’s growth rate at the time of invasion – also known as invasion fitness – assuming that its own influence on the resident’s population dynamics can be neglected at this initial stage.

In our model, the way the resident may exert an influence on the mutant’s fitness is through its impact on the equilibrium frequencies of the (epi)genotypes, hence providing a favorable or unfavorable context for a different genetic architecture. For example, in the two-determinants case, and assuming a stable environment, if the resident population happens to be monomorphic for *α*_*F,r*_ = 1 and thus *α*_*S,r*_ = 0, then *X*_*S*_ reaches its neutral equilibrium distribution (1*/*2; 1*/*2). This means that a mutant with non zero *α*_*S,m*_ will on average have lower fitness than the resident, even though the absolute growth rate, as measured by the Lyapunov exponent approach, would be highest for *α*_*S*_ = 1.

To take such resident / mutant interdependence into account, we computed in the two-determinants case the invasion fitness in all possible pairs of mutant and resident architectures (*α*_*F,m*_, *α*_*F,r*_), with *α*_*F,m*_ and *α*_*F,r*_ taken among 65 equally spread discrete values in the range [0, 1]. Since the environment state fluctuates cyclically, the equilibrium frequencies of the four combinations of possible (epi)genotype values (*X*_*F*_, *X*_*S*_) are periodic functions of time (see Section 2.2 and Appendix A). We thus calculated these frequencies at each of 64 timesteps equally spread over a period *T*. We then calculated, at each timestep and for each pair (*α*_*F,m*_, *α*_*F,r*_), the growth rate of the mutant. The instantaneous growth rate at time *t* of the mutant in a given genetic background can be calculated as:

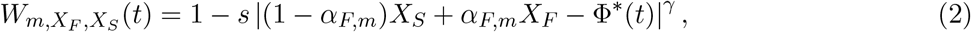

with Φ^*^(*t*) denoting the optimal phenotype at time *t* (either 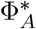 or 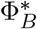). Assuming that the architecture is determined by a modifier locus linked with those controlling the determinants, each combination created at the time of a mutation has its own independent dynamics and long-term relative growth rate, which is not impacted by the change in frequencies in the population and can thus be calculated as the arithmetic mean of the 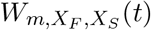 over the period.

The average invasion fitness of a mutant’s architecture depends on its probability to appear in each background, represented by background frequencies in the following equation:

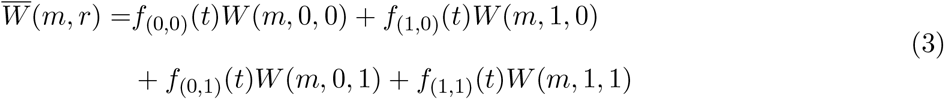

We represent the mutants’ relative invasion fitnesses for our 65×65 pairs (*α*_*m*_, *α*_*r*_) in so-called Pairwise Invasibility Plots (PIP), from which we identify singular strategies and characterize their evolutionary stability (according to Geritz et al. 1998).

### 2.4 Simulations in infinite and finite populations

The numerical simulations for infinite populations are based on the discretisation of the continuous model described below. Although the model is valid for any number of inheritance systems, the numerical results are shown only for two inheritance systems in the case of infinite populations. Then, the population is divided in four groups according to their (epi)genotype: *g*(0,0), *g*(1,0), *g*(0,1) and *g*(1,1). Further, each subpopulation is indexed by individual architecture *α*_*F*_ (simply denoted *α* in the following lines). The density within each subpopulation is driven by

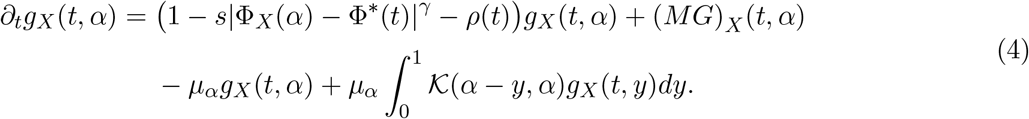

In this equation:

- 1 − *s*|Φ_*X*_(*α*) − Φ^*^(*t*)|^*γ*^ − *ρ*(*t*) is the percapita growth rate, including selection and density dependent saturation where *ρ*(*t*) is the total population density :

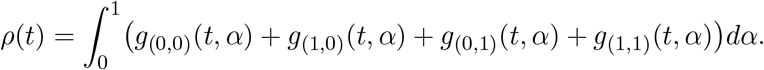
- (*MG*)_*X*_ (*t, α*) describes the (epi)genotype mutation dynamics through the matrix operation *MG* defined by

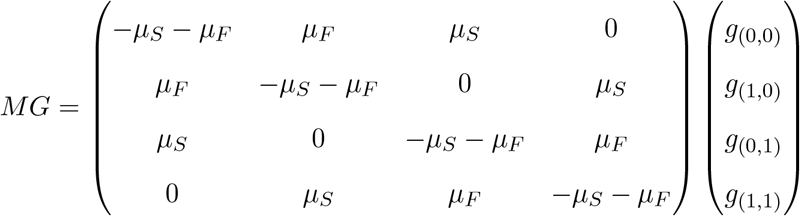
- *μ*_*α*_*g*_*X*_(*t, α*) represents the mutation toward other trait architecture where *μ*_*α*_ is the mutation rate on *α*,
- 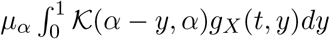 corresponds to the mutation from other trait architecture to the current one; in this term 𝒦 is the mutation kernel that is assumed to be a Gaussian distribution centered on the progenitors trait architecture, conditioned to take values in [0, 1].

The values of *α* are uniformly discretised in [0, 1]. The time discretisation is based on an explicit Runge-Kutta (RK2) scheme to reduce computational time and improve accuracy. The integral terms are computed according to standard quadrature methods.

The model (4) is non-linear due to the density-dependent saturation. Nevertheless, it can be recast into a linear problem, and, in the periodic setting, the Lyapunov exponent *λ*(***α***) coincides with the averaged density over one period 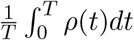 in the case of a monomorphic population when mutations are neglected, see Appendix A.

For finite populations, the stochastic birth and death process described in Section 2.1 is simulated using a Monte-Carlo approach. We opted for a fixed time step discretisation to reduce computational time. Contrary to the infinite population density model (Equation 4), the population remains finite and fluctuates around the carrying capacity *K*. The generalization of the model to any number of inheritance systems is straightforward, at no computational cost, and so the Monte-Carlo approach is suitable when the number of inheritance systems is larger than three.

## 3 Results and discussion

The evolution of the trait architecture ***α*** may stem from two distinct effects: (i) its impact on the average fitness across the two environments, regardless of the various mutation rates and (ii) its control of the phenotypic contribution of inheritance systems with different mutation rates and thus presumably different variational properties. Our focus is on the second effect, that is, on the implications of the mutation rates on the trait architecture. We thus begin our analysis (Section 3.1) by setting *γ* = 1 such that the first effect is removed. Indeed, the averaged fitness does not depend on the phenotype when *γ* = 1, that is, in mathematical terms: 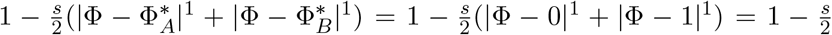 for all Φ.

In Section 3.2, we then investigate the sensitivity of our conclusions to changes in the shape of the trait fitness function.

### 3.1 Architectures evolve in response to the degree of environmental instability

#### 3.1.1 Composite architectures are selected against when the fitness function is linear

To assess which architecture (pure or composite) is optimal under various degrees of environmental instability, we first simulated temporal dynamics in the simple case *I* = 2 (two inheritance systems, fast and slow, with respective mutation rates *μ*_*F*_ and *μ*_*S*_). In this particular case, we recall that respective contributions of the fast and slow inheritance systems are denoted *α*_*F*_ and *α*_*S*_ = 1 − *α*_*F*_. In this first set of simulations, the values of *α*_*F*_ are equidistributed within the population at initial time (polymorphic case). As illustrated in Figure 2, we only observed in the long run selection for one or the other of the two extreme, non-composite trait architectures (*α*_*F*_ = 0 and, symmetrically, *α*_*S*_ = 1, or *α*_*F*_ = 1 and *α*_*S*_ = 0), depending on the environmental cycle duration *T* .

**Figure 2:**
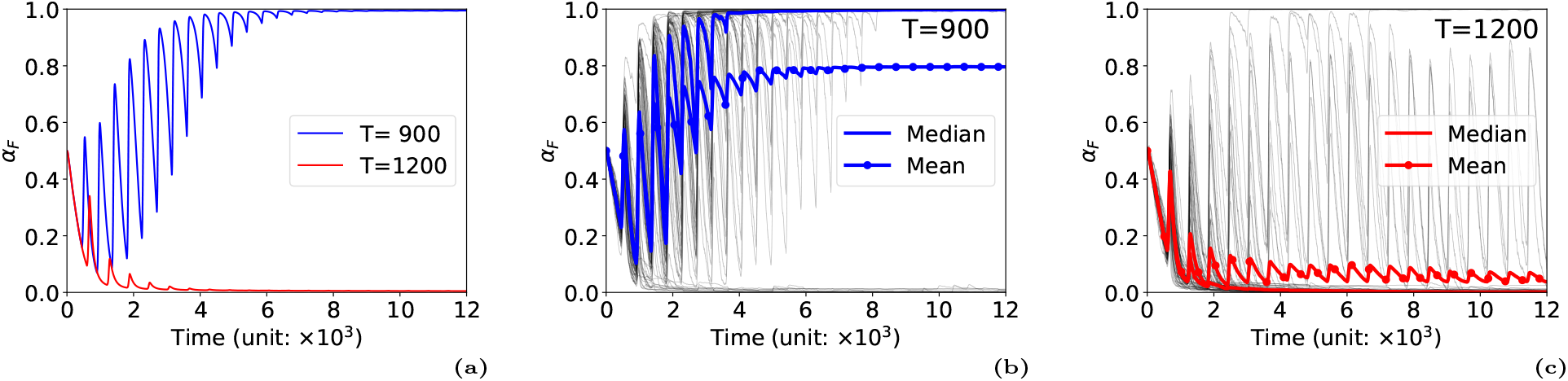
Temporal dynamics of the trait architecture with polymorphic initial configuration. (a) Simulation of the deterministic PDE model (Equation 4), corresponding to an infinite population size, with a polymorphic initial configuration where all *α*_*F*_ values have the same frequency. The mean *α*_*F*_ in the population is plotted over time for two values of the time period of environmental change *T*. We observed the selection towards *α*_*F*_ = 1 for the smaller period *T* = 900 and the selection towards *α*_*F*_ = 0 for the larger period *T* = 1200. (b-c) Monte-Carlo simulations associated with finite population sizes, respectively for *T* = 900 (b) and *T* = 1200 (c). The equilibrium population size is set to *K* = 2^16^ ≈ 65000. The initial value of *α*_*F*_ is drawn uniformly within the population. Each black line represents the trajectory of the averaged value 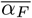 within the population over time. The colored lines represent the mean and median of 64 simulation replicates.

Thus, in this setting, only the slow inheritance system should contribute to the trait (*α*_*F*_ = 0) in scarcely changing environments (large *T* values); on the contrary, in unstable environments, only the fast inheritance system is expected to contribute (*α*_*F*_ = 1). Interestingly, persistent intermediate values of *α*_*F*_ (0 *< α*_*F*_ *<* 1) were never observed. This phenomenon can be illustrated by considering the relative fitnesses of different *α*_*F*_ as a function of *T*, as shown in Figure 3(a) where the intermediate value *α*_*F*_ = 0.5 never has maximal fitness (similar outcomes were observed for other intermediate values 0 *< α*_*F*_ *<* 1, not shown). Notably, at very high frequency of environmental change (small period *T*), fitness differences were found to be negligible.

**Figure 3:**
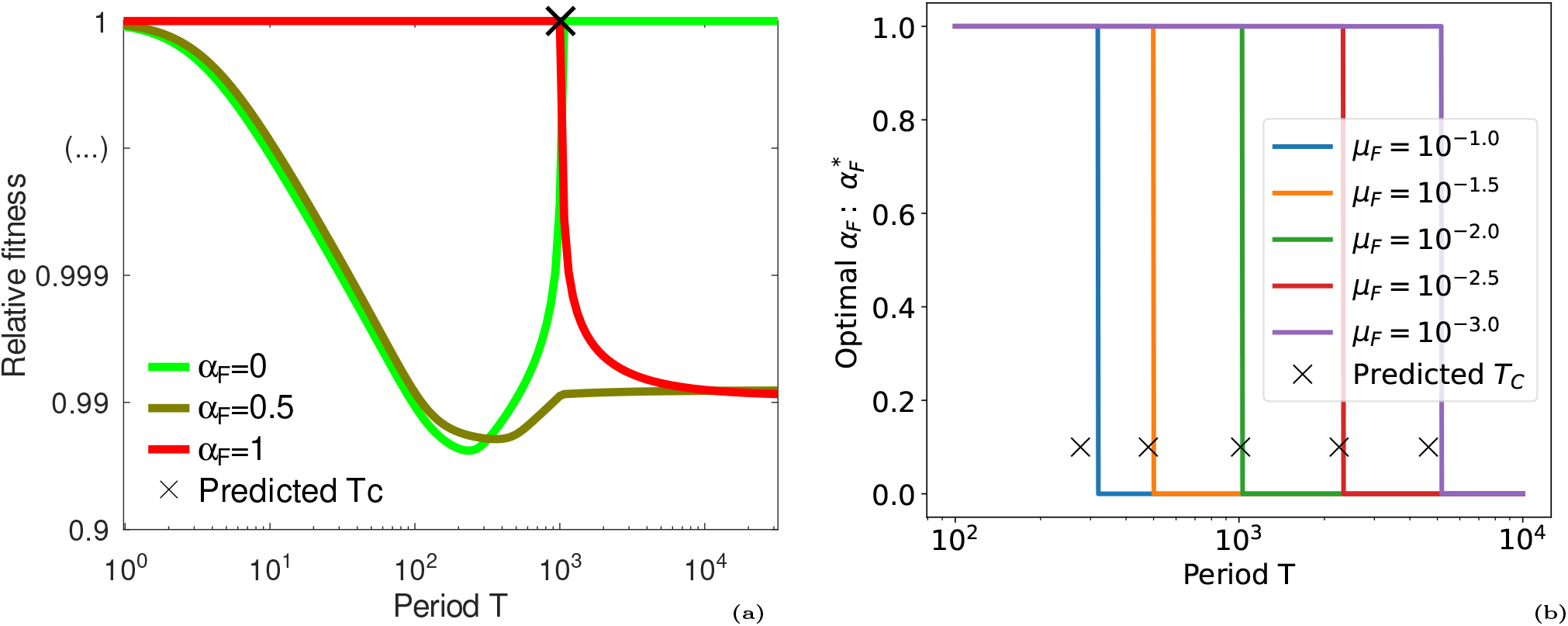
The Lyapunov exponent is maximal for non-composite architectures. (a) Relative fitness 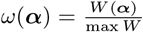 is represented as a function of the period *T*, for three different values of *α*_*F*_ = 0, 0.5, 1. The maximal fitness is always attained at extremal values *α*_*F*_ = 0 or *α*_*F*_ = 1. The analytical value *T*_*c*_ of the transition between ‘unstable’ and ‘stable’ environments is reported by a cross-mark (Equation 5). (b) The optimal value (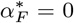 or 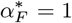) is plotted as a function of the period *T* for different values of the fast mutation rate *μ*_*F*_. Again, the switching time *T*_*c*_ is well approximated by the analytical expression (Equation 5) when *μ*_*F*_ is sufficiently distant from the values of *s* (10^−1^) and *μ*_*S*_ (10^−4^).

In fact, the counter-selection of intermediate *α*_*F*_ values (and symmetrically, of intermediate *α*_*S*_ values) turned out to be a theorem, whose statement is given below:

*Assume that the trait fitness function is linear (γ* = 1*), and I* ≥ 2. *Then the Lyapunov exponent λ*(***α***) *is a convex function of the trait architecture* ***α***.

This theorem is a consequence of the convexity properties of the spectral radius with respect to the diagonal coefficients (see *e*.*g*. Kingman 1961; Cohen 1981). A proof can be found in Appendix B for the sake of completeness. An immediate consequence of this theorem is that optimal values of the trait architecture are reached at some extremal point of the simplex set {***α*** = (*α*_*i*_) : (∀*i*)*α*_*i*_ ≥ 0, and ∑_*i*_ *α*_*i*_ = 1}. Alternatively speaking, composite trait architectures should always be selected against when the selection function is linear.

Another implication of this theorem is the occurrence of sharp transitions, at specific threshold periods, corresponding to switches between optimal trait architectures ***α***^*^. In the case *I* = 2, there is one such switch at a critical value *T* = *T*_*c*_, quantitatively separating unstable environments (*T < T*_*c*_) where the Lyapunov exponent *λ*(***α***) is maximal at *α*_*F*_ = 1 (full contribution of the fast inheritance system), from stable ones (*T > T*_*c*_) where the Lyapunov exponent *λ*(***α***) is maximal at *α*_*F*_ = 0 (full contribution of the slow inheritance medium), see Figure 3(b). The value of *T*_*c*_ can be computed analytically in the regime *μ*_*S*_ ≪ *μ*_*F*_, by seeking the value of *T* such that *λ*(0) = *λ*(1). This was performed, and lead to the following expression (after some further simplifications, see Appendix C for details):

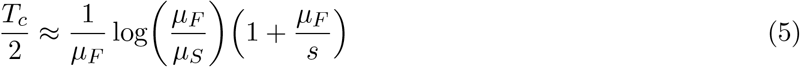

This approximation is valid in the regime *μ*_*S*_ ≪ *μ*_*F*_ ≪ *s*, that is, when the mutation rates are far apart, and much below the maximal fitness defect *s*. It could be further simplified by neglecting 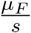, because selection is assumed to be strong enough so that it has limited influence on the threshold period *T*_*c*_.

As mentioned above, non-trivial outcomes are expected in the intermediate regime, when the rate of environmental change falls in the range of mutation rates. Formula (5) makes this expectation more precise as the switch occurs when the duration of one stasis (the half-period 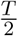) echoes the fast mutation rate, up to a logarithmic correction. The latter correction is essential as it contains the ratio between the two mutation rates (fast versus slow). Interestingly, although selection is obviously essential for this process, its strength *s* only weakly influences the value of the switching period (no dependency at the leading order in formula (5)). With more than two determinants (differing in their mutation rates), several switching periods are expected, as exemplified in the case *I* = 3 (see Appendix D, Figure 7).

#### 3.1.2 Epistasis and contingency in the evolution of trait architectures

A mutant genetic architecture may be more or less successful at invading particular (epi)genetic backgrounds. For example, putting more weight on the fast inheritance system may be deleterious if this system is not in the appropriate state in a certain environment (see Section 2). This creates an evolutionary interaction between a resident genetic architecture and the mutant architecture. Importantly, this is ignored in the Lyapunov exponent approach, where the fitnesses of different architectures are compared after each has reached its own (epi)genetic equilibrium frequencies. In contrast, this is well captured by the adaptive dynamics framework. In this framework, the mutant invasion dynamics is approximated through its growth rate at low frequency within a resident population under deterministic dynamics (infinite population size approximation), that is when it is so rare that it does not itself impact the resident’s dynamics. This growth rate, calculated as detailed in Section 2, is represented in Figure 4(a) for 4225 (65^2^) pairs of mutant and resident strategies.

**Figure 4:**
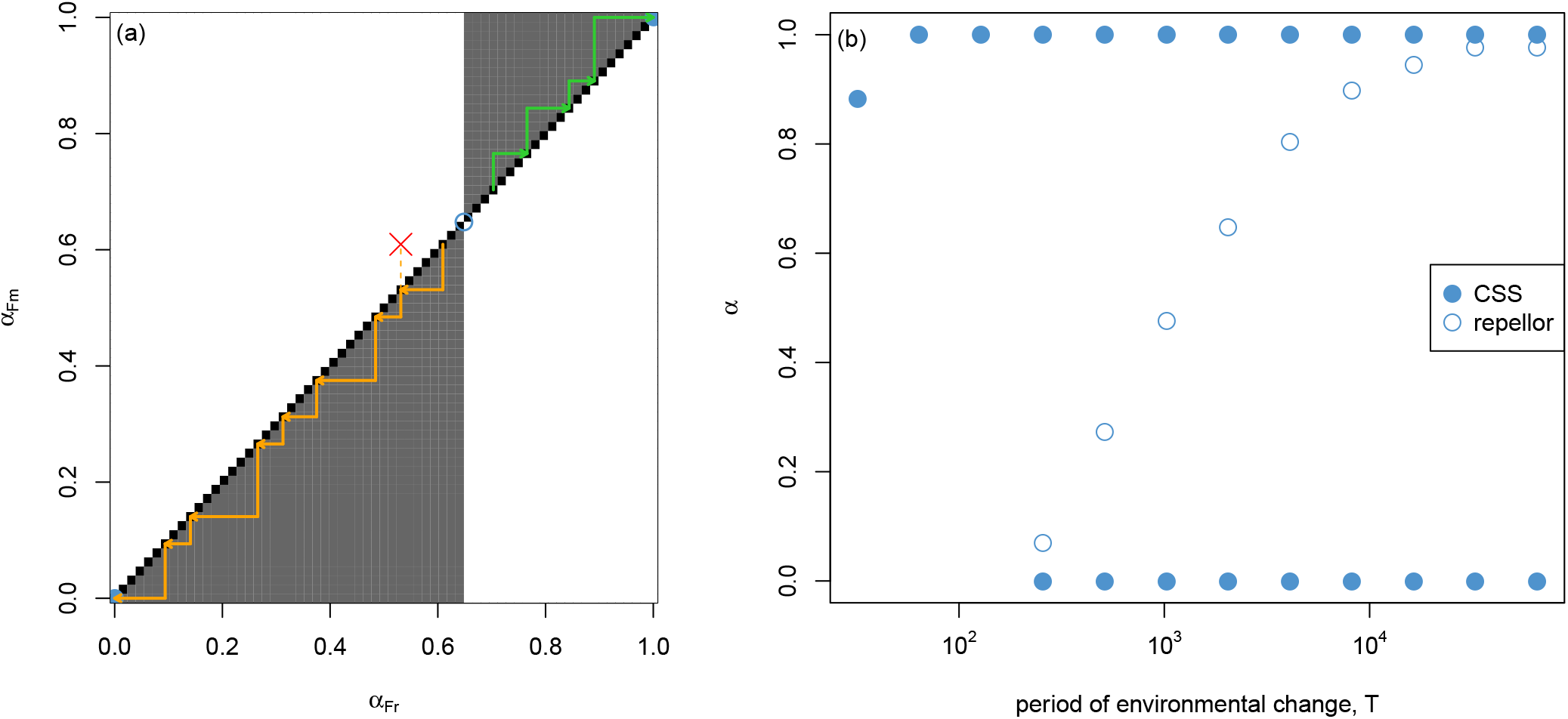
Adaptive dynamics of trait architectures. (a) A pairwise invasibility plot showing how evolutionary trajectories of the genetic architecture may depend on historical contingency. Black squares represent mutant/resident pairs where the relative growth rate of the mutant *λ*(*α*_*Fm*_, *α*_*Fr*_) is close to 1 (*±*10^−9^), while grey and white squares represent pairs where *λ*(*α*_*Fm*_, *α*_*Fr*_) *>* 1 and *<* 1, respectively. A first example evolutionary trajectory is shown in orange: a mutant with a lower contribution of the ‘fast’ component *α*_*Fm*_ appears in a population with *α*_*Fr*_ close to 0.6. The mutant increases in frequency (*i*.*e*. it *invades*; grey square). It is expected to reach fixation (*i*.*e*. replace the resident) because the latter cannot invade back when it is rare (white square at the opposite coordinate, red cross). The evolutionary trajectory continues downwards until the architecture *α*_*F*_ = 0 is fixed. Starting from a slightly different architecture (*α*_*Fr*_ ≈ 0.7, in green), the evolutionary trajectory goes in the opposite direction towards *α*_*F*_ = 1, showing how the evolution of the genetic architecture may be contingent on the ancestral architecture of a population. Mutants increasing *α*_*F*_ in the orange trajectory or decreasing *α*_*F*_ in the blue one, not represented, may have appeared and failed to invade. The evolutionary repellor at *α*_*F*_ ≈ 0.65 is represented by an empty dot and the two CSS attractors at *α*_*F*_ = 0 and 1 by plain dots. Parameter values: *γ* = 1, *T* = 2048, *s* = 0.1. (b) Singular strategies observed under a linear trade-off (*γ* = 1) and different frequencies of environmental change (*x*-axis in log scale). Singular strategies are either repellors (empty dots) or CSS attractors of the evolutionary dynamics (plain dots).

Based on this approach, we identified two types of singular strategies, as represented in Figure 4(a): evolutionary repellors from which evolutionary trajectories tend to move away, and convergent stable strategies (CSS) that represent possible long-term evolutionary attractors. Evolutionary trajectories starting below or above the repellor will be markedly different. If the initial genotype has an architecture *α*_*F*_ below the repellor, *e*.*g*. 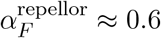 in Figure 4(a), mutants with lower *α*_*F*_ will tend to invade and reach fixation. Because residents with higher *α*_*F*_ consistently fail to invade on average, these evolutionary dynamics will thus lead *α*_*F*_ downwards until *α*_*F*_ = 0 is reached. The reverse occurs for starting points above the repellor. Thus, the existence of a repellor creates a contingency of the evolutionary dynamics on the initial genotype; whether, or rather how frequently it can be overcome by drift will depend on population features (but see Section 3.3) and the distribution of mutation effects on the trait’s architecture. But it should be noted that selection will consistently act against such crossing of the repellor, as shown by the large white areas in Figure 4(a) for which the probability of fixation is lower than that of a neutral mutant.

We next investigate how the types of singular strategies and their distribution change as a function of environmental instability, see Figure 4(b). When the environment is scarcely changing (up to *T* ≈ 256), two CSS attractors (at *α*_*F*_ = 0 and *α*_*F*_ = 1) and one repellor are present and only the position of the repellor changes. In environments that change very rarely, the repellor is very close to *α*_*F*_ = 1, such that one may expect the evolution of architectures towards *α*_*F*_ = 0, that is, with the lowest possible contribution of the most unstable inheritance system. This conclusion is similar to that reached through the Lyapunov exponent approach, and in line with the results of (Rajon and Charlat 2019). Under higher environmental instability, the repellor moves to intermediate values of *α*_*F*_ such that the outcome will depend on the initial genotype. This is again compatible with the results of (Rajon and Charlat 2019) reporting an increase of the average *α*_*F*_ in similar contexts, especially since the initial (epi)genotypes were uniformly distributed between 0 and 1 in their study. Decreasing further the period of environmental change below a threshold of *T* ≈ 256, only the CSS attractor at *α*_*F*_ equal or close to 1 remains.

The aforementioned conclusions derived from the adaptive dynamics were supported by Monte-Carlo numerical simulations initialized with monomorphic populations, that is, where all individuals initially have the same trait architecture: either *α*_*F*_ = 0.2 as in Figure 5(a), or *α*_*F*_ = 0.8 as in Figure 5(b). For sufficiently large population size, *K* = 4096, we observed evolution towards one or the other of the two extreme, non-composite trait architectures (blue trajectories), in agreement with results of Figure 4(a). Individual trajectories are depicted respectively in Figure 5(e) and Figure 5(h), showing a clear selection of the extreme values, with only few trajectories crossing the repellor in Figure 5(e), and none in Figure 5(h) among 256 replicates. The other panels (smaller population sizes) will be discussed below in Section 3.3.

**Figure 5:**
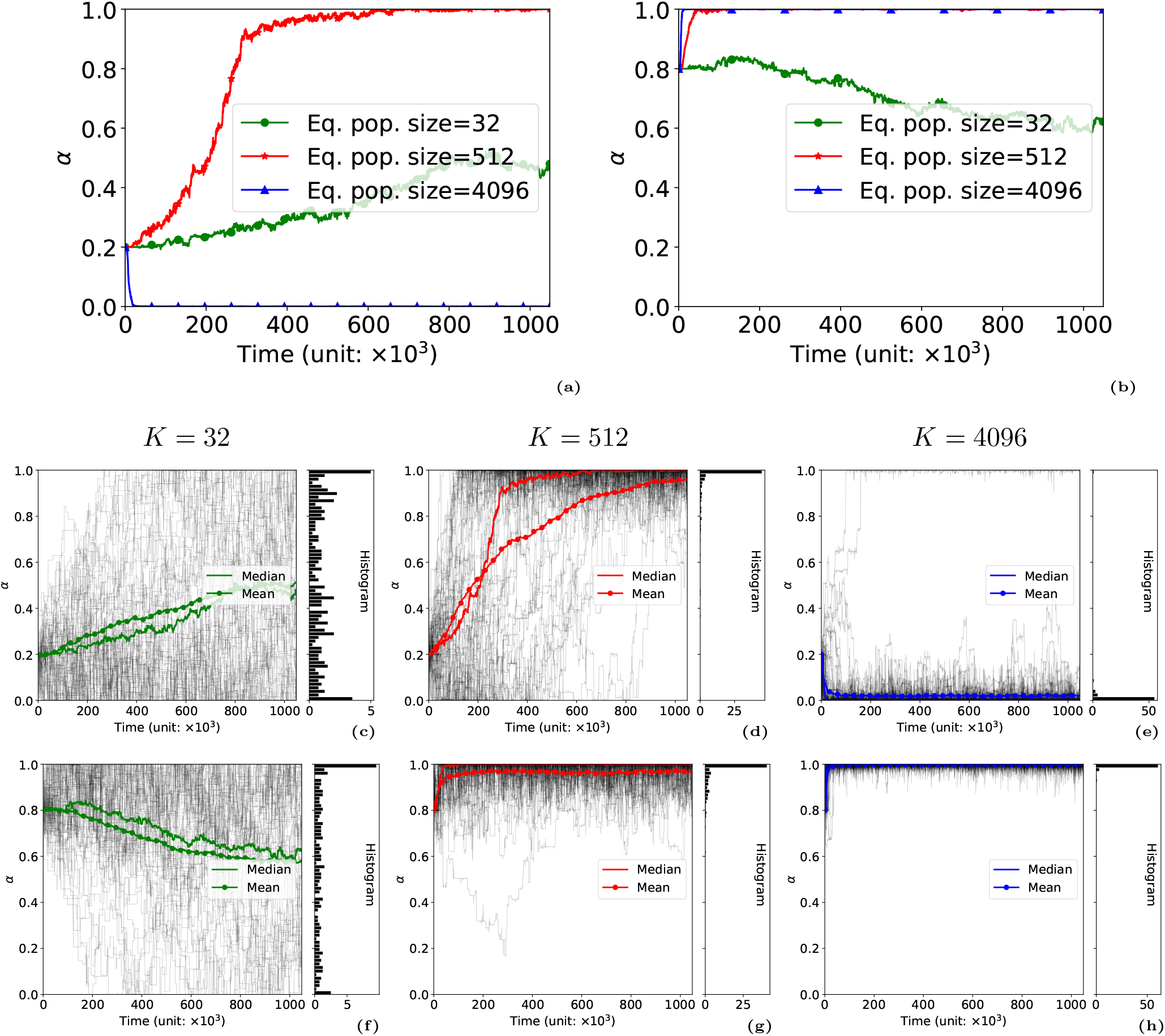
Temporal dynamics of the trait architecture with monomorphic initial configuration. (a-b) Median of the mean value 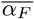 over the population for 256 simulation replicates starting respectively from: *α*_*F*_ = 0.2 (a), and *α*_*F*_ = 0.8 (b), associated with various population size. The (epi)genotypes are initialized with the monomorphic equilibrium distribution, depending on the value of *α*_*F*_. The figure legend indicates the typical population size for each of the three numerical experiments (resp. *K* = 32, 512, 4096). The case of small population size (*K* = 32) exhibits a pattern of neutral evolution, dominated by genetic drift. The cases of intermediate population size (*K* = 512) and large population size (*K* = 4096) exhibit the same trend when starting from *α*_*F*_ = 0.8 (above the repellor value ≈ 0.65), but opposite trends when starting from *α*_*F*_ = 0.2 (below the repellor value). (c-h) Sample of the simulation replicates (*N* = 64): trajectories of the mean value 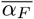 in each simulation out of the sample is plotted in dark line, and the final distribution is plotted as a side histogram. Values are as follows (typical population size *K*, and initial value of the monomorphic trait architecture *α*_*F*_) : *K* = 32, *α*_*F*_ = 0.2 (c); *K* = 512, *α*_*F*_ = 0.2 (d); *K* = 4096; *α*_*F*_ = 0.2 (e); *K* = 32, *α*_*F*_ = 0.8 (f); *K* = 512, *α*_*F*_ = 0.8 (g); *K* = 4096; *α*_*F*_ = 0.8 (h). Parameter values: period T = 2048, s = 0.1 (same as Figure 4).

### 3.2 Sensitivity to the fitness function shape

#### 3.2.1 Composite architectures are selected against when the fitness function is convex

The results of Section 3.1.1 can be extended to the case of a convex fitness function, in the sense of Figure 1 (*γ <* 1). Indeed, the trait architecture fitness inherits convexity from the trait fitness function, as stated in the following theorem.

*Assume that the trait fitness function is convex, and I* ≥ 2. *Then, the Lyapunov exponent λ*(***α***) *is a convex function of the trait architecture* ***α***.

As a by-product of convexity, we deduce that, in the case *I* = 2, the critical time *T* = *T*_*c*_ does not depend on the shape parameter *γ*, provided that *γ* ≤ 1 (convex selection), simply because it is characterized by the Lyapunov exponents of the extreme trait architectures through the equality *λ*(0) = *λ*(1). Since the intensity of selection does not depend on *γ* at the extremal architectures (either 1 or 1 − *s*, and vice-versa), the balance *λ*(0) = *λ*(1) does not involve the parameter *γ*.

It is important to notice that there is no corresponding statement regarding concavity, that is, the trait architecture fitness does not inherit concavity from the trait fitness function in general. For instance, the selection function in the case *γ* = 1 (Section 3.1.1) is both convex and concave because it is linear. However, the trait fitness architecture is strictly convex in this case. It is the transition between several (epi)genetic states by mutation, and the environmental variation which generates such strict convexity.

Since the convexity property is a strong feature, which holds true even in the case of a linear selection function, we expect that convexity of the Lyapunov exponent persists beyond the linear regime, even if the trait fitness function is not convex anymore, say slightly concave. Robustness of the previous conclusions with respect to the shape of the fitness function is the purpose of the next section.

#### 3.2.2 Concave fitness functions may select for composite architectures

Contrary to convexity, concave fitness functions may result in positive selection for a balanced contribution of various determinants – that is 0 *< α*_*F*_ *<* 1 – compare Figure 6(a) with Figure 2(a). Thus, in order to assess the robustness of our main conclusions from Section 3.1.1, we investigated the optimal trait architecture maximizing the Lyapunov exponent *λ*(***α***) in the case of a concave selection function (*γ* ≥ 1), see Figure 6(b). Interestingly, the concept of a critical switching value *T*_*c*_ still exists when composite trait architectures are selected for, even for large values of *γ*. Fast inheritance systems have more weight in the selected architecture 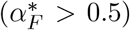 when *T < T*_*c*_. In contrast, slow inheritance systems have more weight in the selected architecture 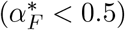, when *T > T*_*c*_. Moreover, the transition is sharp at *T* = *T*_*c*_ (switch between a clear balance on the side of fast inheritance towards a clear balance on the opposite side), a feature shared with the convex case, compare Figure 6(b) with Figure 3(b). Although we lack a formula for the critical value *T*_*c*_ in the concave case, we see clearly that it does not depend on the shape parameter (*γ*). Although this may appear surprising at first glance, it might be expected from the fact that the critical value does not depend on the shape parameter (*γ*), nor on the intensity of selection (*s*) at the leading order in the convex case, see Equation (5).

**Figure 6:**
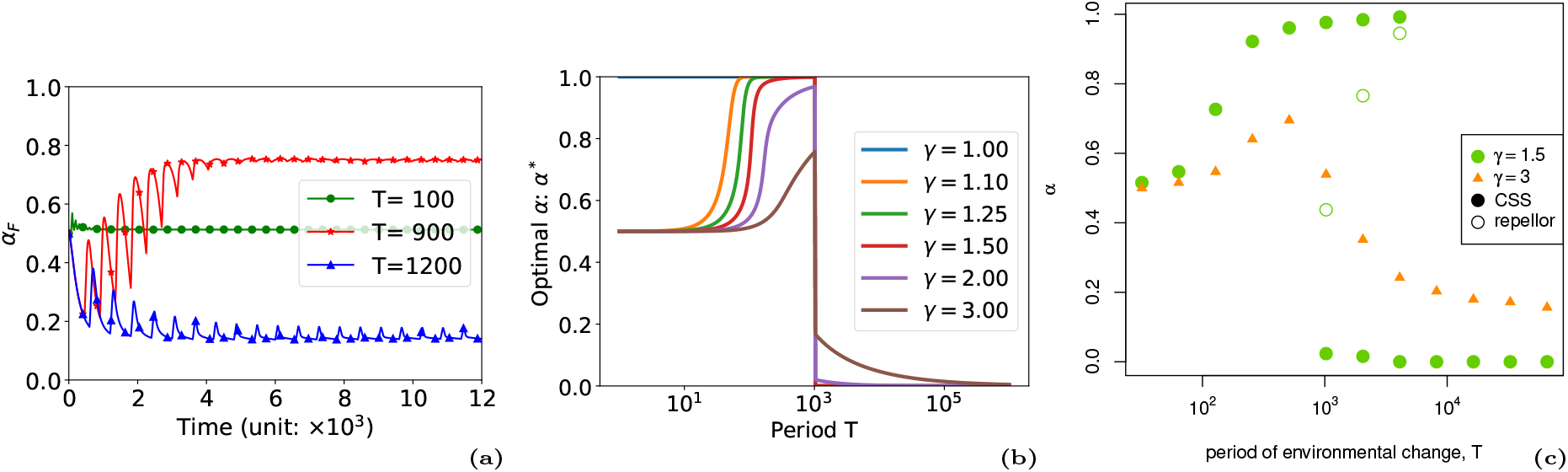
Main outcomes for concave fitness functions. (a) Temporal dynamics for the deterministic PDE models of infinite population size for three values of the time period *T* = 100, *T* = 900, and *T* = 1200, and *γ* = 3 (analogous to Figure 2(a)). The mean value of *α*_*F*_ in the population is plotted over time. In contrast with the linear case, we observed the selection for non-composite architectures, especially when the environment is very unstable (*T* = 100). (b) The optimal value 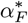 is represented as a function of the shape parameter *γ >* 1 (analogous to Figure 3(b)). In contrast with the linear case, non-composite architectures can result in a higher relative fitness *ω*, especially when the environment is very unstable, or when *γ* is large. (c) Singular strategies observed under a concave selection function (*γ >* 1) and different periods of environmental change). Singular strategies are either repellors (empty dots) or CSS attractors of the evolutionary dynamics (plain dots or triangles).

Notably, non-composite architectures can still be selected for, for moderate shape parameter (*γ* ≲ 1.5) and period *T* close enough to the switching period *T*_*c*_. For large *γ*, the fastest inheritance system 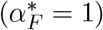 is never selected for, and the slowest one is selected for only in very stable environments (*T* ≫ *T*_*c*_).

Regarding the adaptive dynamics, the results in Figure 6(c) are consistent with the previous results. When the selection function is moderately concave (here, *γ* = 1.5), and the environment changes infrequently (*T >* 8192), a single CSS attractor is present at *α*_*F*_ close to 0. For intermediate regimes of environmental variation, a repellor and the two attractors at respectively low and high *α*_*F*_ are present, similar to the case where *γ* = 1 (compare with Figure 6(b)). As environmental variations become more frequent, at *T <* 1024 a single CSS remains and the expected selected architecture tends to intermediate values of *α*_*F*_, ending up at *α*_*F*_ = 0.5 for very frequent changes. As concavity increases to *γ* = 3, the transition between an architecture with a large contribution of the determinant with the lowest mutation rate (low *α*_*F*_) in scarcely varying environments to an architecture with a higher *α*_*F*_ no longer includes a repellor, nor a very large contribution of the least stable determinant – *i*.*e. α*_*F*_ does not reach values close to 1.

### 3.3 Sensitivity to population size: a new case of sign inversion

We used simulations to assess the predictions of the adaptive dynamics approach under more or less effective selective regimes, that is, under various populations sizes. Simulations behave as expected when the population size is sufficiently large (*K* = 4096), that is *α*_*F*_ goes to 0 when the population is initiated with *α*_*F*_ below the repellor and to 1 otherwise. In very small populations, selection is overall inefficient and *α*_*F*_ tends to become uniformly distributed, independently of its initial value (*K* = 32 in Figure 5). This is also expected since genetic drift tends to dominate over selection in these conditions. But in populations of intermediate sizes (*K* = 512 in Figure 5), simulations reveal a complex pattern. When the initial value *α*_*F*_ = 0.8 is above the repellor, it evolves toward 1, as expected. Yet, surprisingly, *α*_*F*_ also evolves toward 1 when it is initiated *below* the repellor, at initial value *α*_*F*_ = 0.2. In other words, we found that not only the efficiency of selection, but also its direction, depends on the population size (compare panels (d) and (e) of Figure 5, where the only parameter that differs is *K*).

We reached the following interpretation for this surprising pattern. The evolution of *α*_*F*_ in fluctuating environments is dynamically driven by two opposite selective pressures: (1) upon shifts to a new environment, high values of *α*_*F*_ are selected for, because these give more weight to the “fast” inheritance system, that has a higher chance of being in a state that matches the new environment, or to reach this state through an (epi)mutation; (2) during periods of environmental stasis, low values of *α*_*F*_ are selected for, because these reduce the cost of frequent deleterious (epi)mutations. Depending on the frequency of environmental change, one of these processes tends to dominate, provided that populations are large enough. A plausible interpretation for the observed pattern (that is, for a deterministic evolution of *α*_*F*_ toward high values in populations of “intermediate size”) relies on noting that the strength and duration of the selection associated with the two processes differ. As a result, one process, subject to stronger selection than the other, may become effective in populations where the other is not. More specifically, what we see here is in our view stemming for a strong (but infrequent) selection toward high *α*_*F*_ values upon environmental change and a weak (but durable) selection toward low *α*_*F*_ values during periods of environmental stasis. Both are effective in large populations, but weak and durable selection toward low *α*_*F*_ dominates. In contrast, in populations of intermediate size, only the strong albeit infrequent selection toward high *α*_*F*_ operates.

With this explanation in mind, we realized that other authors have recently shed light on a very similar process that they coined ‘sign inversion’ (Raynes, Wylie, et al. 2018; Raynes, Burch, and Weinreich 2021). This expression appropriately emphasizes the main point of the explanation: changes in population size can not only make selection more or less efficient, but may also inverse its direction in systems where two opposing pressures are at play and differ in their intensity and duration. In large populations, both pressures are effective, but weak and long lasting selection may dominate; under a certain threshold population size, only the strong but infrequent selection is effective. Following the discovery of this process in a study focusing on the evolution of mutation rates, Raynes, Wylie, et al. 2018 have noted that this process may be far more than anecdotal, because many traits can be subject to opposing selective pressures, for example because of temporal variations in the environment. Our study provides one more example of situations where this process would take place.

## 4 Conclusion

Our analysis was aimed at assessing the hypothesis that heterogeneity in the mutation rates of a trait’s determinants (of whatever nature) may affect the evolution of its architecture, that is, of the respective contributions of the different determinants. Our main conclusion is that in a variety of conditions, in both stable and unstable environments, non-composite architectures are effectively selected for, meaning that trait values would tend to depend upon a single determinant, or several determinants characterized by similar mutation rates. This sheds light on prior simulation results (Rajon and Charlat 2019) as we now understand how the degree of environmental variation conditions which determinant, rather than which mixture of determinants, takes part to the trait architecture. This focus on the mutational portfolio of traits contributes to the general understanding of the evolution of a trait’s architecture, as a complement to the more common focus on the number of contributing genes, their epistatic and pleiotropic properties, and the distribution of their respective effects.

Let us emphasize that a non-composite trait architecture may still be “complex” in the sense of involving many determinants (*e*.*g*. Ungerer et al. 2002; Flint and Mackay 2009; Kemper, Visscher, and Goddard 2012; Yengo et al. 2022). For example, Rajon and Plotkin 2013, in a model assuming a single mutation rate, have shown that many genes should contribute to traits subject to a stabilizing selection of mild intensity. In such a context, where the environment is kept stable, our model predicts that the architecture should be non-composite. Overall, a complex yet non composite architecture would then be expected, involving the most stable determinants.

The impact of environmental heterogeneity on the evolution of trait architecture has also been the subject of previous studies. Yeaman and Whitlock 2011 have shown that spatial differences in traits optima leads the evolution of recombination landscapes toward simplified architecture where clusters of genes have large impact of the phenotype. Using yet another framework, the results of (Hansen et al. 2006) suggest that directional selection, producing an ever going requirement for adaptation should result in an increase in mutational effects through changes in epistatic interactions, provided that mutations are rare enough.

In our study, we focused on temporal environmental fluctuations on the evolution of the trait architecture. More specifically, we assumed that environmental changes reverse the direction of selection, with fitness gains modeled as concave, linear, or strictly convex functions. These shapes were found critical: only a concave fitness function resulted in composite architectures, provided that the degree of instability overcomes a threshold. The same conclusion holds for a mixture of concave and convex fitness shapes, like the typical case of a Gaussian function centered on a moving optimum, see Appendix D, Figure 8(a). In such a case, the amplitude of the fluctuations may be critical because Gaussian functions are concave close to the center, but convex at the border, as can be seen in Appendix D, Figure 8(b).

**Figure 7:**
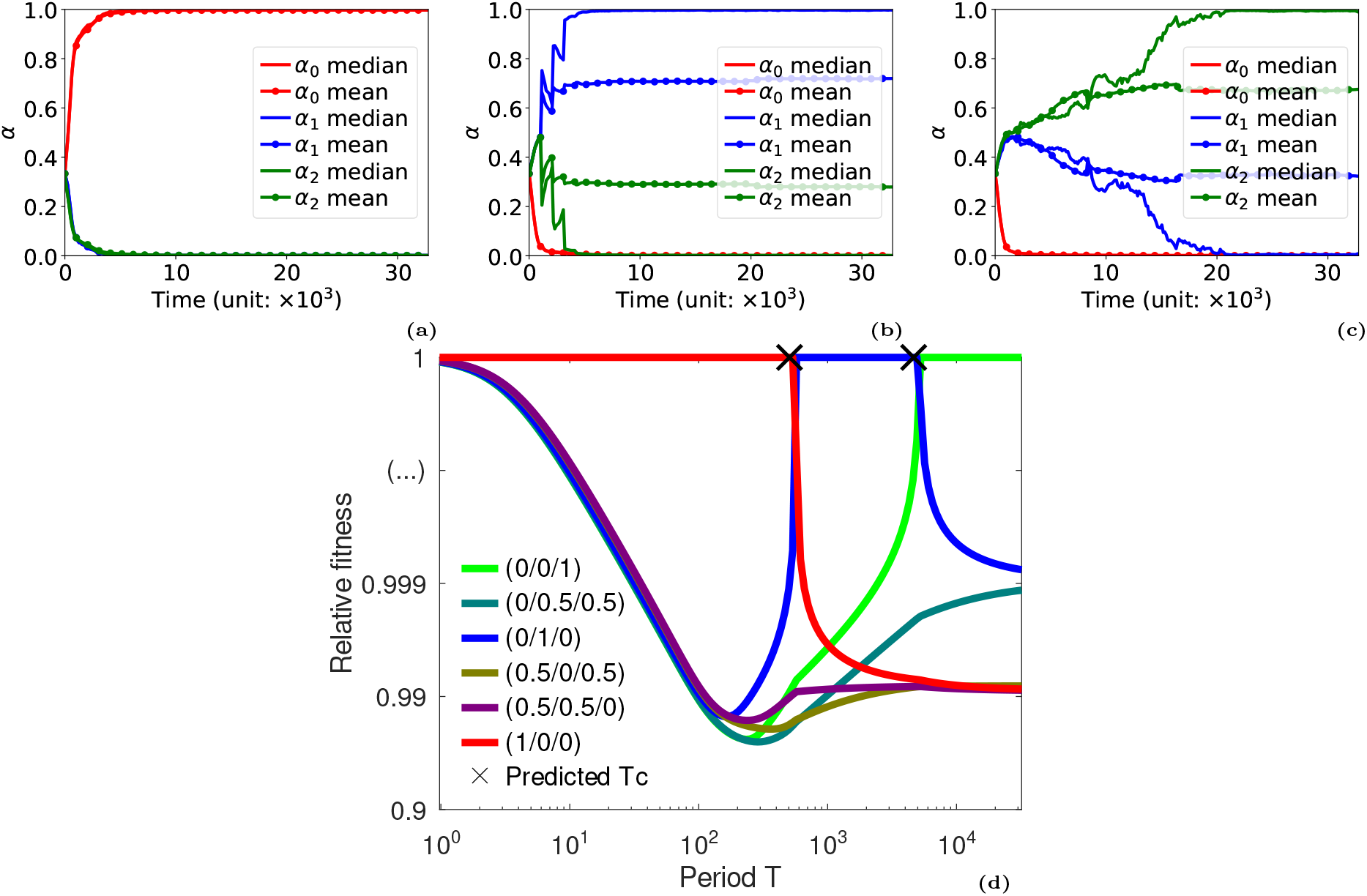
Results for three media *I* = 3 and a linear selection function. *γ* = 1. (a-b-c) Temporal dynamics for the Monte-Carlo simulations with three determinants (*I* = 3) associated with mutation rates *μ*_*i*_ = 10^−2^, 10^−3^, 10^−4^ (analogous to Figure 2(b-c)). We observed the selection towards full contribution of one of the three determinant for each value of the time period among *T* = 128 (a: fast switching environment, selection of the largest mutation rate), *T* = 2048 (b: intermediate switching environment, selection of the intermediate mutation rate), *T* = 16384 (c: slow switching environment, selection of the slowest mutation rate) (d) Relative fitness 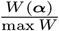 as a function of the period *T*, for different values of the trait architecture ***α*** (analogous to Figure 3(a)). The maximal fitness is always attained at extremal values of the trait architecture (full contribution of a single determinant).

**Figure 8:**
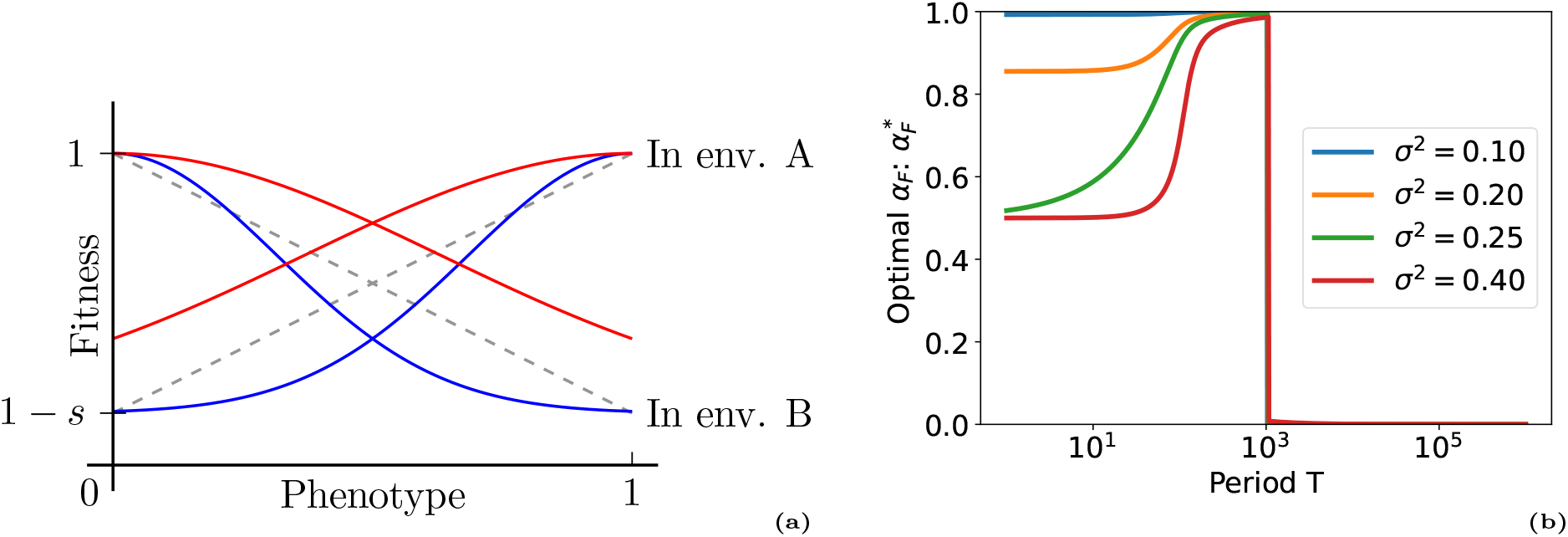
Gaussian fitness function and associated optimal trait architecture: (a) Shape of the fitness function for Gaussian selection (ie. mixed concave and convex shape): 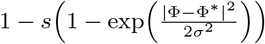, for *σ*^2^ = 0.1 (plain blue) and *σ*^2^ = 0.4 (plain red). (b) The optimal value 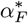 is represented as a function of the shape parameter *σ*^2^ (analogous to Figure 3(b) and Figure 6(b)). Only the nearly concave fitness function (*σ*^2^ ≳ 0.20) resulted in composite architectures, provided that the degree of instability overcomes the usual threshold (*T < T*_*c*_).

While the shape of fitness functions acting on the trait is a key parameter, it will only be relevant if selection on the architecture is effective, which ultimately depends on its strength, in relation with population size. We developed complementary approaches to investigate the strength of selection on the trait architecture. The Lyapunov exponent approach revealed conditions where composite architecture are not optimal, but it missed the notion that the evolution of trait architecture is driven by the background variations, accumulated by different determinants through their distinct mutation rates and across environmental fluctuations. The adaptive dynamics approach captured this effect, by assessing the average strength of selection over an environmental fluctuation period. But Monte Carlo simulations in populations of finite size revealed a singular aspect of these dynamics, related to the effect of drift. Simulations in large populations were in agreement with the adaptive dynamics predictions, and those in very small populations were as expected driven by drift only. Yet, in populations of intermediate sizes, we found that the trait architecture evolves deterministically in the opposite direction as in large populations. We interpret this result as a new occurrence of ‘sign inversion’ (Raynes, Wylie, et al. 2018; Raynes, Burch, and Weinreich 2021), stemming from the periodic alternation between long time windows of weak selection for a heavy contribution of stable inheritance systems (inefficient in populations of intermediate sizes) and short time windows of strong selection for a heavy contribution of unstable systems. This result further complexifies our understanding of the evolution of genetic architectures, arguing in favor of a systematic use of population genetics models and, as an aside, for further mathematical analyses that would clarify and generalize the conditions for the occurrence of sign inversion.

## Appendices

### A. Model reduction for Lyapunov exponent computation

In this section, we aim to detail the links between the continuous model presented in Section 2.4 and formalized by Equation (4) and the Lyapunov exponent computation. Indeed, the dynamic is driven by the leading eigenvalues in a linear ordinary differential equation system. The presence of the integral term for the trait architecture mutation and the logistic growth rate to model the density dependent saturation make our model non-linear. However, in the case of a monomorphic population, when the trait architecture mutations are neglected, we demonstrate a correspondance with the Floquet spectral theory of periodic linear systems, such that the average of the density over one period, 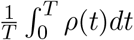, and the Lyapunov exponent defined in Equation (1), coincide.

Assume that mutations on the trait architecture *α* are negligible. For a given trait architecture *α* and group *X* ∈ {(0, 0), (1, 0), (0, 1), (1, 1)} the equation (4) becomes

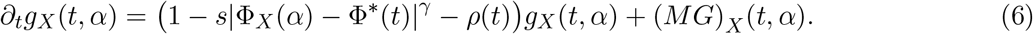

Depending on the optimal phenotype Φ^*^, the system gathering equations (6) writes

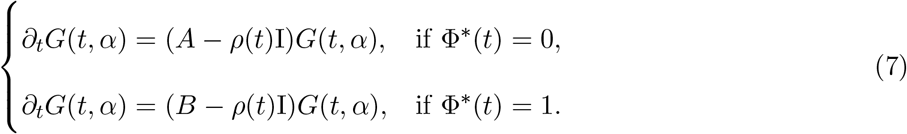

Assume that both matrix *A* and *B* can be diagonalised. Thus, there exist a constant invertible matrix *P*_*A*_ (resp. *P*_*B*_) and a diagonal matrix *D*_*A*_ (resp. *D*_*B*_) such that 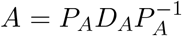 (resp. 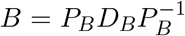). So the systems (7) becomes

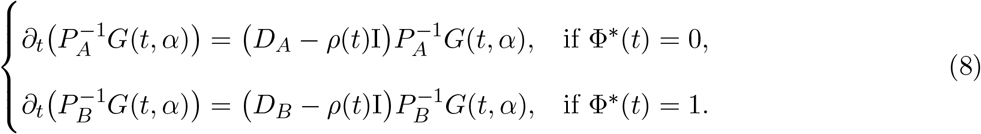

With the notations 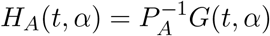 and 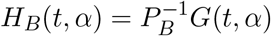 this systems writes

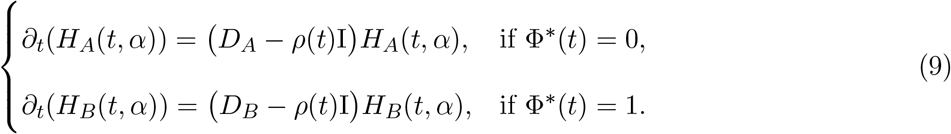

#### Relations between *H* components

Let *H*_*A,i*_ denotes the *i*^*th*^ components of *H*_*A*_ and *d*_*A,i*_ the *i*^*th*^ diagonal term of *D*_*A*_. Assume that the *H*_*A,i*_ never vanishes, then

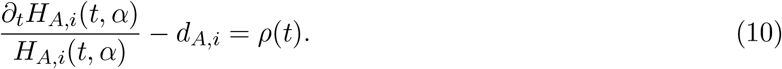

Thus, for any pair (*i, j*)

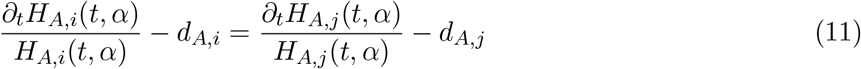

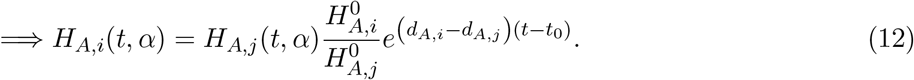

with 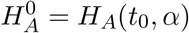. Therefore, the relative dynamics of the components of the vector *H* depend on the difference between the eigenvalues of *A*. By the same reasoning and using similar notations it also comes

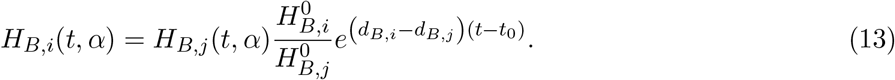

#### Solution characterization using total population size

According to systems (9), as long as Φ^*^(*t*) remains constant on the time interval [*t*_0_, *t*] with *t > t*_0_ the solutions are

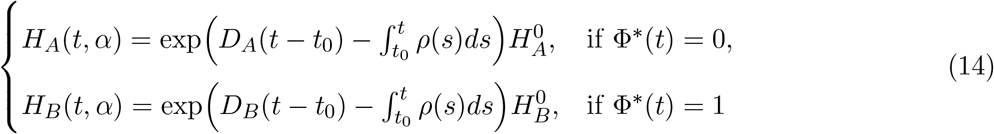

with 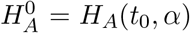 and 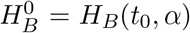. Note that these formulas rely on 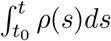 which is not explicitly known. Using the relation *G*(*t, α*) = *P*_*A*_*H*_*A*_(*t, α*) and *G*(*t, α*) = *P*_*B*_*H*_*B*_(*t, α*) it comes

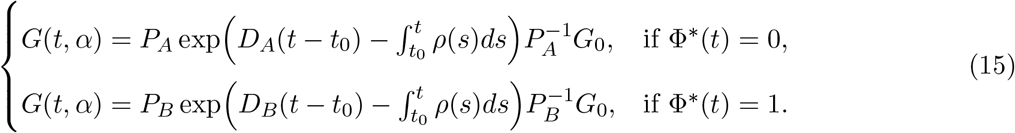

where *G*_0_ = (*t*_0_, *α*). These formulas simply into

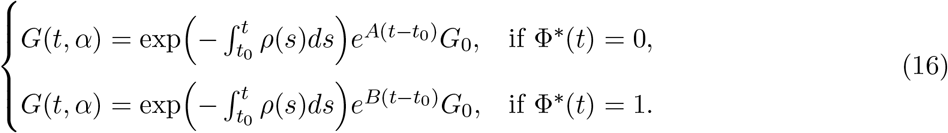

##### A.1 Groups dynamic in stable environment

Without loss of generality, let’s consider the case of Φ^*^(*t*) = 0 and drop the subscript *A* to shorten notations. Using *G*(*t, α*) = *PH*(*t, α*) one can deduce that the expression of *G*_*i*_ the *i*^*th*^ components of *G* is

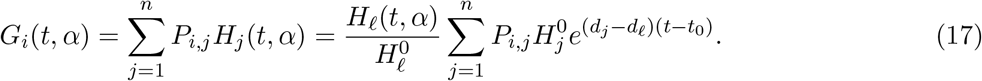

where *ℓ* is an arbitrary index to be fixed later. Thus, the total population size is

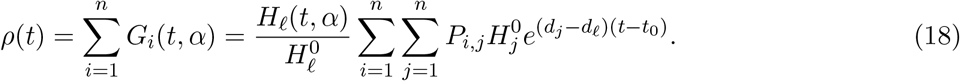

Therefore, the relative group size within the total population size is

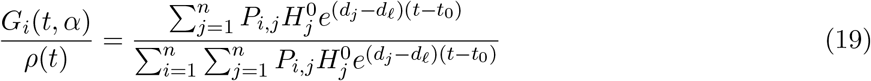

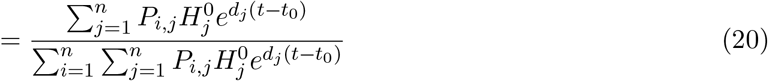

Thus, the relative group size dynamic is fully determined by the eigenvalues of matrix *A*. Moreover, this shows that the whole dynamic is driven by the *A* matrix leading eigenvalue:

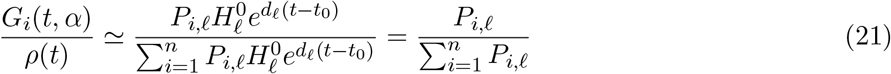

assuming that *d*_*ℓ*_ is the leading eigenvalue. Note that 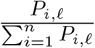 correspond to the normalised value of the *i*^*th*^ component of the eigenvector associated to *d*_*ℓ*_. That is, the normalised eigenvector associated with the largest eigenvalue of *A* approximates the relative group sizes.

##### A.2 Groups dynamic in periodic environment

Let’s consider a periodic environment of period *T*. Without loose of generality let assume that Φ^*^(*t*) = 0 for 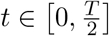 and Φ^*^(*t*) = 1 for 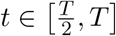. Thus, starting from *G*_0_ = *G*(0, *α*) and using formulas (16) the solution at *t* = *T* is

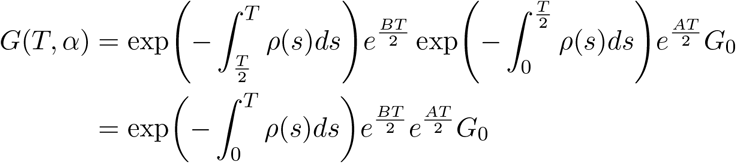

Assume that the solution becomes periodic, namely *G*(*T, α*) = *G*_0_ then the previous formula lead to

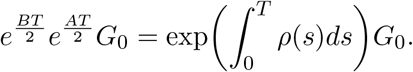

So *G* is the eigenvector associated to the leading eigenvalue 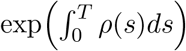 of 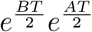. Moreover, this demonstrates that the leading eigenvalue is real.

Up to renormalization, the eigenvector *G*_0_ can be interpreted as the relative group sizes within the whole population. Indeed, the relative size of a group *ℓ* is simply 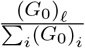.

### B. Proof of convexity

This section is devoted to the mathematical proof of our main convexity result, that is the convexity of the Lyapunov exponent *λ*(***α***) in the case of a convex selection function (*γ* ≤ 1). This result can be recast in the framework of convexity of dominant eigenvalues of non-negativity matrices. To apply such theory, one should be careful about the convex combination of coefficients during convex interpolation. Indeed, the diagonal coefficients and the off-diagonal coefficients do not play the same role, for as they are not subject to the same constraints (non-negativity constraints off the diagonal). More precisely, convex combinations are arithmetical on the diagonal, but geometrical off the diagonal (see the *θ*−interpolation below). The situation is facilitated in our model as the selection acts only on the diagonal terms via a trait-dependent mortality rate. Nevertheless, we shall present a more general result to emphasize this discrepancy between diagonal and off-diagonal coefficients.

We claim that the following Theorem encompasses our main convexity result:

*Let A, B two matrices with positive entries off the diagonal*^1^, *that is* ∀(*i, j*) *a*_*ij*_ *>* 0, *b*_*ij*_ *>* 0.

*Let θ* ∈ (0, 1). *Consider the matrix C*_*θ*_ *defined by the following term-by-term interpolation:*

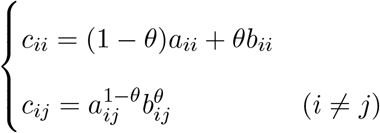

*Then the dominant (Perron) eigenvalues λ*(*A*), *λ*(*B*), *λ*(*C*_*θ*_) *satisfy the following convex inequality:*

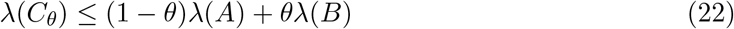

We make the following comments:

- This result is classically credited to (Kingman 1961).
- The result can be extended straightforwardly to irreducible matrices by a limiting argument.
- We are interested in the particular case where *A* and *B* differ only by their diagonal entries, that is, *a*_*ij*_ = *b*_*ij*_ if *i* ≠ *j*. In this case, the interpolation 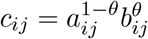 is trivial, and the convex inequality (22) can be reformulated as: *λ*((1 − *θ*)*A* + *θB*) ≤ (1 − *θ*)*λ*(*A*) + *θλ*(*B*). Alternatively speaking, the dominant eigenvalue is convex with respect to its diagonal.
- In fact, the proof can be extended to the Floquet eigenvalues in the periodic setting, which is precisely our focus:

*Let A*(*t*), *B*(*t*) *a pair of time-dependent, periodic, matrices with positive entries off the diagonal. Let θ* ∈ (0, 1). *Consider the matrix C*_*θ*_(*t*) *defined pointwise as follows:*

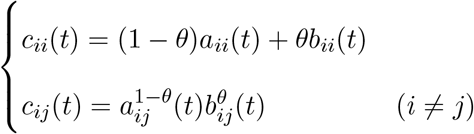

*Then the dominant (Floquet) eigenvalues λ*(*A*), *λ*(*B*), *λ*(*C*_*θ*_) *satisfy the following convex inequality:*

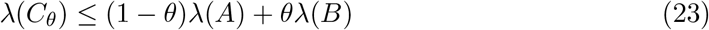

We present first the proof of the static case (Perron eigenvalues), then we extend the method to the time-periodic setting (Floquet eigenvalues).

Our approach is based on (Clairambault, Gaubert, and Lepoutre 2011), see also (Cohen 1981).

#### The static case

We need the following eigen-elements: *X* (resp. *Y*) the positive (right) eigenvector of *A* (resp. *B*), and Φ the positive (left) eigenvector of *C*. We have accordingly:

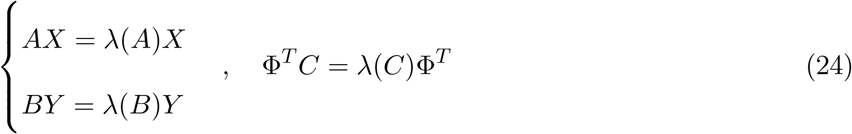

We define *Z* by the pointwise interpolation: 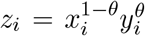. Then, we evaluate the Collatz-Wielandt

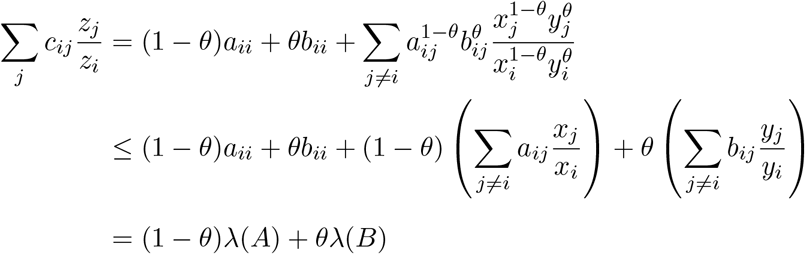

where we have used the Hölder inequality in the second line, and the fact that *X* and *Y* are eigenvectors in the last line. We deduce that

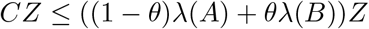

Finally, multiplying by Φ^*T*^, we find:

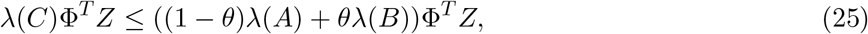

hence, *λ*(*C*) ≤ (1 − *θ*)*λ*(*A*) + *θλ*(*B*).

#### The time-periodic case

The proof is almost identical, but the fact that we have to consider time-periodic eigenvectors *X*(*t*) (resp. *Y* (*t*)) and Φ(*t*) such that:

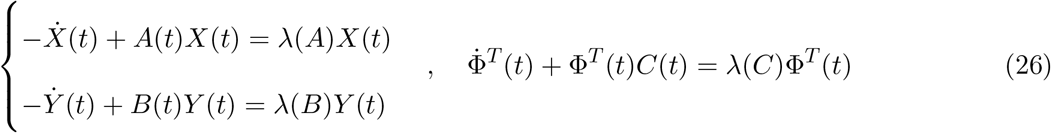

We define again *Z*(*t*) pointwise: 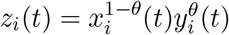. Then, we evaluate the Collatz-Wielandt ratio:

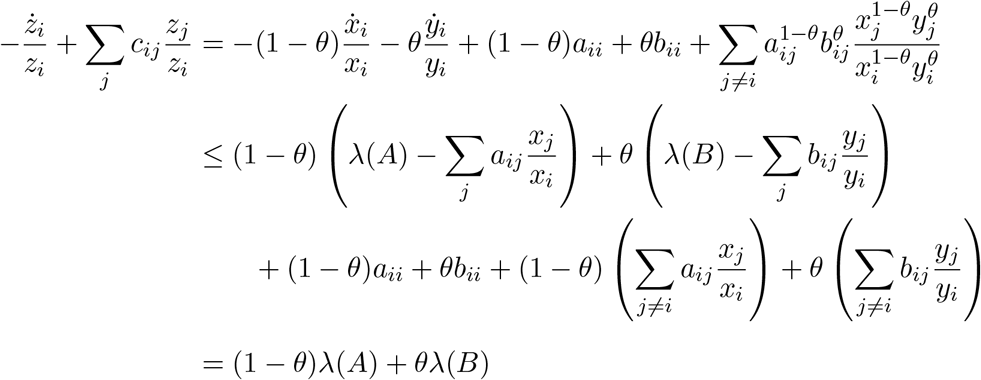

We deduce that

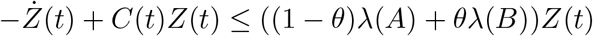

To conclude, we multiply by *ϕ*^*T*^ and integrate other one period:

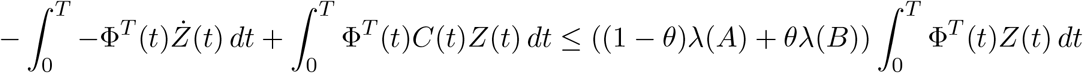

Integrating by parts the first term (using periodicity), we find

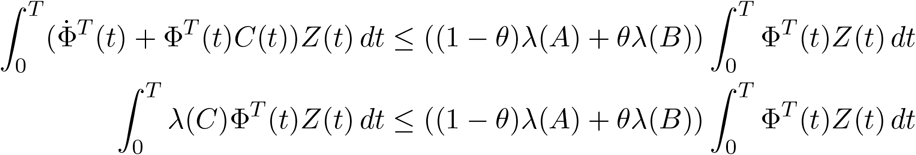

Hence, *λ*(*C*) ≤ (1 − *θ*)*λ*(*A*) + *θλ*(*B*).

### C Computation of the critical time *T*_*c*_

This section presents the analytical computation of the critical time *T*_*c*_ of the transition between ‘unstable’ and ‘stable’. According to our main results, see Section 3.1.1: *when the trait fitness function is linear (γ* = 1*), then the Lyapunov exponent λ*(***α***) *is a convex function of the trait architecture* ***α***. Consequently, for the two determinant case the optimal trait architecture is either^2^ *α*_*F*_ = 0, or *α*_*F*_ = 1 and the transition occurs when the Lyapunov exponents satisfy *λ*(0) = *λ*(1). Thus, the critical time *T*_*c*_ can be computed by solving this equation. However, according to Equation (1), the Lyapunov exponent is the largest eigenvalue of a product of matrix exponential. Thus, its computation is very tedious so we relied on Maple (2022) to perform the preliminary steps of the calculation.

As mentioned above, the goal is to solve *λ*(0) = *λ*(1) thus the first step consists in determining the Lyapunov exponent for *α*_*F*_ = 0 and *α*_*F*_ = 1. To shorten the notation, we abbreviate *α*_*F*_ to *α*. According to Maple, the eigenvalues of 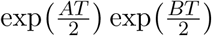 when *α* = 0 are

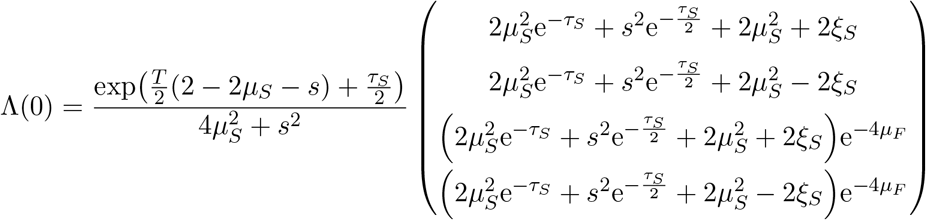

with

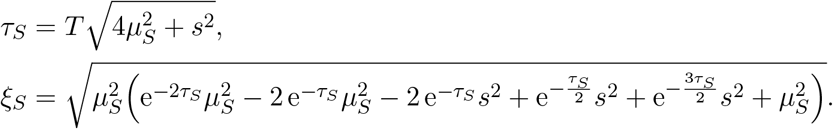

The mutation rate *μ*_*F*_ is strictly positive so 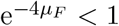 and thus the largest eigenvalue is either Λ(0)_1_ or Λ(0)_2_. Besides, *ξ*_*S*_ simplifies into

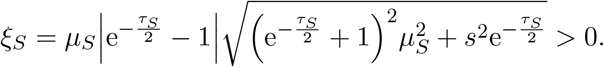

Thus, when *α* = 0 the largest eigenvalue is

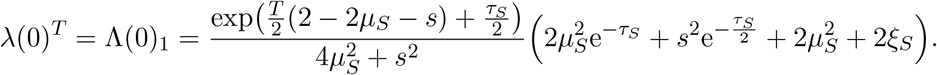

Similarly, using Maple, we found that the eigenvalues of 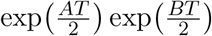 when *α* = 1 are

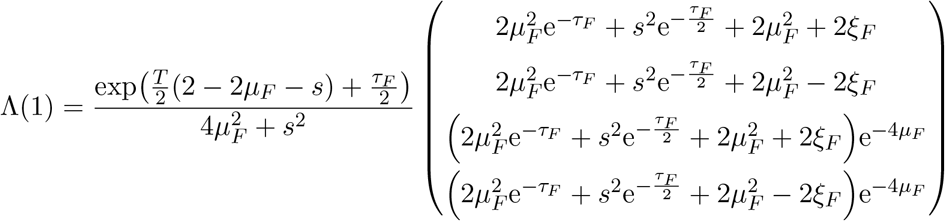

with

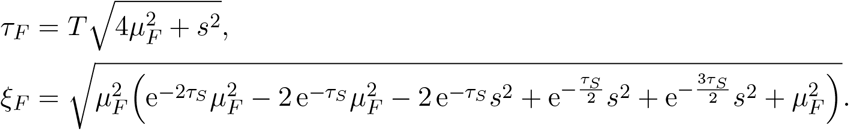

As previously *ξ*_*F*_ simplifies into

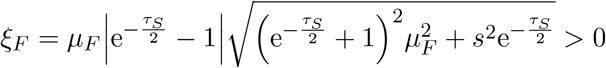

and using the same arguments we found that when *α* = 1 the largest eigenvalue is

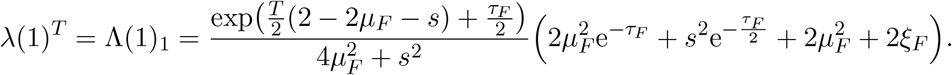

Therefore, the critical time *T*_*c*_ can be computed by solving Λ(0)_1_ = Λ(1)_1_. In practice, we were unable to solve this equation. Even the numerical approximation of the solution requires particular care due to the presence of large terms in the exponentials. Nevertheless, the terms of the form 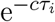 with *c >* 0 a constant and *i* ∈ {*F, S*} are negligible in the regime *μ*_*F*_ ≪ *s*. Thus, for *α* = 0 it comes 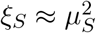 and then

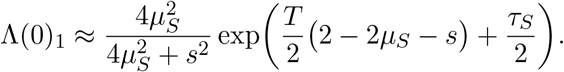

Similarly, for *α* = 0 it comes 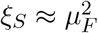 and then

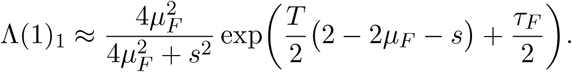

So, the approximated critical time *T*_*c*_ is the solution of the equation:

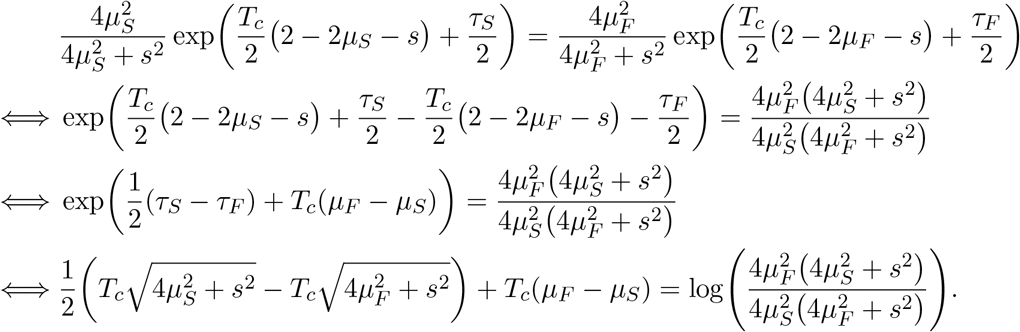

Thus, the critical time is approximately

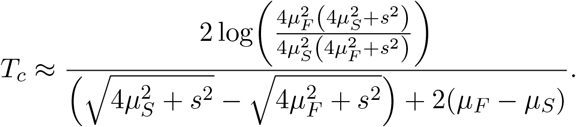

However, as it stands, the contribution of each parameter to the critical time remains complex. Thus, according to Taylor expansions in the regime *μ*_*S*_ ≪ *μ*_*F*_ ≪ *s*, this formula simplifies into

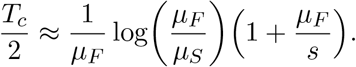

### D Supplementary figures

## Footnotes

1 usually referred to as a Metzler matrix

2 We recall that in the two determinant case ***α*** = (*α*_*F*_, *α*_*S*_) with *α*_*S*_ = 1 − *α*_*F*_ .

## References

Andre, J. B. and B. Godelle. “The evolution of mutation rate in finite asexual populations”. Genetics 172 (2006).

Clairambault, J., S. Gaubert, and T. Lepoutre. “Circadian rhythm and cell population growth”. Mathematical and Computer Modelling 53 (2011).

Cohen, J. E. “Convexity of the dominant eigenvalue of an essentially nonnegative matrix”. Proceedings of the American Mathematical Society 81 (1981).

Danchin, E. “Avatars of information: towards an inclusive evolutionary synthesis”. Trends Ecol Evol 28 (2013).

Denkena, J., F. Johannes, and M. Colomé-Tatché. “Region-level epimutation rates in Arabidopsis thaliana”. Heredity 127 (2021).

Dodson, A. E. and J. Rine. “Heritable capture of heterochromatin dynamics in Saccharomyces cerevisiae”. eLife 4 (2015).

Flint, J. and T. F. C. Mackay. “Genetic architecture of quantitative traits in mice, flies, and humans”. Genome Res. 19 (2009).

Geritz, S., E. Kisdi, G. Meszéna, and J. Metz. “Evolutionarily singular strategies and the adaptive growth and branching of the evolutionary tree”. Evol Ecol 12 (1998).

Graaf, A. van der, R. Wardenaar, D. A. Neumann, et al. “Rate, spectrum, and evolutionary dynamics of spontaneous epimutations”. Proceedings of the National Academy of Sciences 112 (2015).

Hansen, T. F., J. M. Alvarez-Castro, A. J. R. Carter, and J. Hermiss. “Evolution of Genetic Architecture under Directional Selection”. Evolution 60 (2006).

Hodgkinson, A. and A. Eyre-Walker. “Variation in the mutation rate across mammalian genomes”. Nature Reviews Genetics 12 (2011).

Ishii, K., H. Matsuda, Y. Iwasa, and A. Sasaki. “Evolutionarily stable mutation rate in a periodically changing environment”. Genetics 121 (1989).

Johannes, F., E. Porcher, F. K. Teixeira, et al. “Assessing the impact of transgenerational epigenetic variation on complex traits”. PLoS Genet 5 (2009).

Johnson, T. “Beneficial mutations, hitchhiking and the evolution of mutation rates in sexual populations”. Genetics 151 (1999).

Kemper, K. E., P. M. Visscher, and M. E. Goddard. “Genetic architecture of body size in mammals”. Genome Biol. 13 (2012).

Kingman, J. F. C. “A convexity property of positive matrices”. The Quarterly Journal of Mathematics 12 (1961).

Metz, J., R. Nisbet, and S. Geritz. “How should we define ‘fitness’ for general ecological scenarios?” Trends Ecol Evol 7 (1992).

Oman, M., A. Alam, and R. W. Ness. “How Sequence Context-Dependent Mutability Drives Mutation Rate Variation in the Genome”. Genome Biology and Evolution 14 (2022).

Rajon, E. and J. B. Plotkin. “The evolution of genetic architectures underlying quantitative traits”. Proc. R. Soc. Lond. B (2013).

Rajon, E. and S. Charlat. “(In)exhaustible Suppliers for Evolution? Epistatic Selection Tunes the Adaptive Potential of Nongenetic Inheritance”. The American Naturalist 194 (2019).

Rando, O. J. and K. J. Verstrepen. “Timescales of genetic and epigenetic inheritance”. Cell 128 (2007).

Raynes, Y., C. L. Burch, and D. M. Weinreich. When good mutations go bad: how population size can change the direction of natural selection. preprint. Evolutionary Biology, 2021.

Raynes, Y., C. S. Wylie, P. D. Sniegowski, and D. M. Weinreich. “Sign of selection on mutation rate modifiers depends on population size”. Proceedings of the National Academy of Sciences 115 (2018).

Rechavi, O., G. Minevich, and O. Hobert. “Transgenerational Inheritance of an Acquired Small RNA-Based Antiviral Response in C. elegans”. Cell 147 (2011).

Ungerer, M. C., S. S. Halldorsdottir, J. L. Modliszewski, T. F. C. Mackay, and M. D. Purugganan. “Quantitative trait loci for inflorescence development in Arabidopsis thaliana”. Genetics 160 (2002).

Yeaman, S. and M. C. Whitlock. “The genetic architecture of adaptation under migration-selection balance”. Evolution 65 (2011).

Yengo, L., S. Vedantam, E. Marouli, et al. “A saturated map of common genetic variants associated with human height”. Nature (2022).

